# Regression dynamic causal modeling for resting-state fMRI

**DOI:** 10.1101/2020.08.12.247536

**Authors:** Stefan Frässle, Samuel J. Harrison, Jakob Heinzle, Brett A. Clementz, Carol A. Tamminga, John A. Sweeney, Elliot S. Gershon, Matcheri S. Keshavan, Godfrey D. Pearlson, Albert Powers, Klaas E. Stephan

**Author notes:** to whom correspondence should be addressed **Correspondence:** Stefan Frässle, University of Zurich & ETH Zurich, Translational Neuromodeling Unit (TNU), Institute for Biomedical Engineering, Wilfriedstrasse 6, 8032 Zurich, Switzerland.

## Abstract

“Resting-state” functional magnetic resonance imaging (rs-fMRI) is widely used to study brain connectivity. So far, researchers have been restricted to measures of functional connectivity that are computationally efficient but undirected, or to effective connectivity estimates that are directed but limited to small networks.

Here, we show that a method recently developed for task-fMRI – regression dynamic causal modeling (rDCM) – extends to rs-fMRI and offers both directional estimates and scalability to whole-brain networks. First, simulations demonstrate that rDCM faithfully recovers parameter values over a wide range of signal-to-noise ratios and repetition times. Second, we test construct validity of rDCM in relation to an established model of effective connectivity, spectral DCM. Using rs-fMRI data from nearly 200 healthy participants, rDCM produces biologically plausible results consistent with estimates by spectral DCM. Importantly, rDCM is computationally highly efficient, reconstructing whole-brain networks (>200 areas) within minutes on standard hardware. This opens promising new avenues for connectomics.

## 1. Introduction

“Resting-state” functional magnetic resonance imaging (rs-fMRI) has long been used to examine the functional organization of the brain during unconstrained cognition when no specific experimental manipulations are present (Biswal et al., 1995; Biswal et al., 1997). Specifically, resting-state fMRI has revealed spontaneous (endogenous) fluctuations in the blood oxygen level dependent (BOLD) signal that are highly structured across the brain (Fox et al., 2005; Greicius et al., 2009; Raichle et al., 2001). Over the last two decades, rs-fMRI has become one of the most vibrant fields in neuroimaging (for reviews, see Smith et al., 2013; van den Heuvel and Hulshoff Pol, 2010) and now plays a central role in disciplines such as connectomics (Craddock et al., 2013) and network neuroscience (Bassett and Sporns, 2017). This is partly due to its practical simplicity, which renders rs-fMRI amenable to subject populations that may otherwise struggle to adhere to complex cognitive tasks, such as neuropsychiatric patients, elderly people, or infants.

So far, rs-fMRI data has been analyzed predominantly in terms of functional connectivity, which represents statistical interdependencies amongst BOLD signal time series from spatially distinct regions of the brain (Friston, 2011). The simplest way to assess functional connectivity is by computing Pearson’s correlation coefficient between the respective BOLD signal time series. Other measures of functional connectivity include partial correlation, coherence, mutual information and independent component analysis (for a comprehensive review, see Karahanoglu and Van De Ville, 2017).

Regardless of the exact approach, these statistical techniques have revealed a number of large-scale networks of correlated temporal patterns in the resting brain. These *resting-state networks (RSN)* are characterized by fairly coherent time courses across its set of inherent components, whereas time courses are sufficiently distinct between networks (Beckmann et al., 2005; Fox and Raichle, 2007; Smith et al., 2009). The most prominent RSN is arguably the default mode network (DMN) which is also frequently referred to as the task-negative network (Raichle et al., 2001; Shulman et al., 1997). The DMN was initially discovered as a consistent set of brain regions that showed deactivations across various attention-demanding, goal-oriented, non-self-referential tasks (for a comprehensive review, see Raichle, 2015). By now, it is clear that the DMN can also be active in specific goal-oriented paradigms such as autobiographical tasks (Spreng, 2012), thinking about others and oneself, or remembering the past and planning the future (Buckner et al., 2008). Since the discovery of the DMN, similar patterns of resting-state coherence have been identified in most cortical systems, giving rise to other RSNs such as the dorsal attention network (DAN; Corbetta and Shulman, 2002; Fox et al., 2006), salience network (SAN; Dosenbach et al., 2007; Menon and Uddin, 2010), and central executive network (CEN; Spreng et al., 2013; 2010). All RSNs persist even during sleep and under various forms of anaesthesia (Picchioni et al., 2013; but see Tagliazucchi and Laufs, 2014), and show high heritability (Fornito et al., 2011; Glahn et al., 2010), suggesting that they might represent fundamental organizing principles of the human brain.

Importantly, mapping the macroscopic functional connectome from rs-fMRI data has not only shed light on the organizational principles in the healthy human brain, but also in disease. Aberrant resting-state functional connectivity has been observed in most psychiatric and neurological disorders (Baker et al., 2019; Buckholtz and Meyer-Lindenberg, 2012; Bullmore and Sporns, 2009; Fornito et al., 2015; Stam, 2014). For example, psychiatric diseases including schizophrenia (Baker et al., 2014; Friston et al., 2016; Lui et al., 2015), depression (Greicius et al., 2007; Wang et al., 2012a) and autism (Courchesne et al., 2007; Hahamy et al., 2015) have all been associated with pathological alterations in resting-state functional connectivity.

Despite these profound contributions to our understanding of the organizing principles of the human brain, measures of functional connectivity essentially describe statistical properties of the data and do not reveal how the data were caused or generated. As such, functional connectivity does not directly capture mechanisms at a latent level of neuronal interactions. Furthermore, measures of functional connectivity are undirected and thus do not capture asymmetries in reciprocal connections. This is problematic because asymmetries in the coupling amongst brain regions have been repeatedly found both in terms of anatomy (Felleman and Van Essen, 1991; Markov et al., 2014a) and function (Frässle et al., 2016; Gazzaniga, 2000; Stephan et al., 2007; Zeki and Shipp, 1988).

In contrast, effective connectivity refers to directed interactions amongst brain regions and can be assessed by exploiting a generative model of the latent (hidden) neuronal states and how these give rise to the observed measurements (Friston, 2011). One of the most widely used generative modeling frameworks for inferring effective connectivity from fMRI data is dynamic causal modeling (DCM; Friston et al., 2003), and variants of DCM have been established to model the resting state, including stochastic DCM (Daunizeau et al., 2009; Li et al., 2011) and spectral DCM (Friston et al., 2014a). While capable of more mechanistic accounts of functional integration during the resting state, these models are restricted to relatively small networks due to computational limitations.

We recently introduced a novel variant of DCM for fMRI – termed regression dynamic causal modeling (rDCM; Frässle et al., 2018a; 2017) – that differs from previous DCMs in several aspects. Most important, rDCM is computationally highly efficient and scales gracefully to very large networks including hundreds of nodes, paving the way for whole-brain effective connectivity analyses (Frässle et al., 2020a). Furthermore, the model can exploit structural connectivity information to constrain inference on directed functional interactions or, where no such information is available, infer optimally sparse representations of whole-brain connectivity patterns. While rDCM was initially designed to work with experimentally controlled perturbations (i.e., task data), the current implementation can, in principle, also by applied to rs-fMRI data. However, this has not been tested thoroughly so far. Here, we assess the face validity and construct validity of rDCM for inferring effective connectivity during the resting state. To this end, we conducted comprehensive simulation analyses and compared rDCM against spectral DCM as an alternative and well-established generative model of rs-fMRI data for small networks. This comparison made use of a large empirical resting-state fMRI dataset acquired by the Bipolar-Schizophrenia Network on Intermediate Phenotypes (B-SNIP) consortium (Tamminga et al., 2013). Finally, we showed that rDCM gracefully scales to large networks with more than 200 areas and enables fast inference on whole-brain effective connectivity patterns from rs-fMRI data at the time-scale of minutes.

## 2 Methods and Materials

### 2.1 Regression dynamic causal modeling

#### 2.1.1 General overview

Regression DCM (rDCM) is a novel variant of DCM for fMRI that enables effective connectivity analyses in large (whole-brain) networks (Frässle et al., 2018a; 2017). This can be achieved by applying several modifications and simplifications to the original DCM framework. In brief, these include (i) translating state and observation equations from time to frequency domain using the Fourier transformation (under stationarity assumptions), (ii) replacing the nonlinear biophysical model of hemodynamics with a linear hemodynamic response function (HRF), (iii) applying a mean field approximation across regions (i.e., connectivity parameters targeting different regions are assumed to be independent), and (iv) specifying conjugate priors on neuronal (i.e., connectivity and driving input) parameters and noise precision. These modifications essentially reformulate the linear DCM in the time domain as a Bayesian linear regression in the frequency domain, resulting in the following likelihood function:

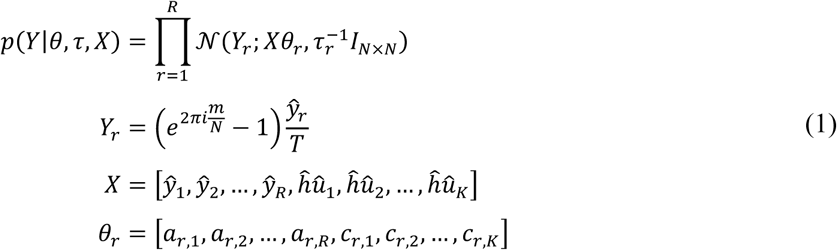

Here, *Y*_*r*_ is the dependent variable in region *r* that is explained as a linear mixture of afferent connections from other regions and direct (driving) inputs. Specifically, *Y*_*r*_ is the Fourier transformation of the temporal derivative of the measured signal in region *r*. Furthermore, *Y*_*r*_ represents the measured BOLD signal in region *r, X* is the design matrix (comprising a set of regressors and explanatory variables), *u*_*k*_ is the k^th^ experimental input, and the hat symbol denotes the discrete Fourier transform (DFT). Additionally, θ_*r*_ represents the parameter vector comprising all connections *a*_*r*,1_, …, *a*_*r,R*_ and all driving input parameters *c*_*r*,1_, …, *c*_*r,K*_ targeting region *r*. Finally, τ_*r*_ denotes the noise precision parameter for region *r* and *I*_*N*×*N*_ is the identity matrix (where *N* denotes the number of data points). Choosing appropriate priors on the parameters and hyperparameters in Eq. (1) (see Frässle et al., 2017) results in a generative model that can be used for inference on the directed connection strengths and inputs.

Under this formulation, inference can be done very efficiently by (iteratively) executing a set of analytical Variational Bayes (VB) update equations concerning the sufficient statistics of the posterior density. In addition, one can derive an expression for the negative (variational) free energy (Friston et al., 2007). The negative free energy represents a lower-bound approximation to the log model evidence that accounts for both model accuracy and complexity. Hence, the negative free energy offers a sensible metric for scoring model goodness and thus serves as a criterion for comparing competing hypotheses (Bishop, 2006). We have recently further augmented rDCM by introducing sparsity constraints into the likelihood of the model in order to allow automatic pruning of fully connected network structures (Frässle et al., 2018a). Notably, the present paper focuses on the original implementation of rDCM without sparsity constraints. A comprehensive description of the generative model of rDCM, including the mathematical details of the neuronal state equation underlying the likelihood function in Eq. (1), can be found elsewhere (Frässle et al., 2018a; 2017).

#### 2.1.2 Application to resting state data

While rDCM was originally designed to work with experimentally controlled perturbations (i.e., task data), it is also possible to fit the model to rs-fMRI data. This can be achieved by “switching off” driving inputs (i.e., setting all input parameters *c*_*i,j*_ to zero in the absence of experimental inputs *u*_*k*_) and, in order to explain activity in any given region, relying on measured data (in the Fourier domain) in regions from which afferent connections are received; compare the likelihood function in Eq. (1). This means that, in contrast to stochastic variants of DCM, the generative model does not explicitly represent endogenous fluctuations in neuronal activity and has no concept of stochastic “innovations” or similar ways noise can drive neuronal activity intrinsically. Instead, endogenous fluctuations of BOLD signal in any given region are predicted as a linear mixture of intrinsic fluctuations from other regions.

In this study, we examine the face validity and construct validity of rDCM in its specific application to rs-fMRI data. Concerning face validity, we investigate whether the particular formulation of rDCM allows for veridical inference on effective connectivity, using simulated BOLD data fluctuations that were generated by means of a deterministic generative model with stochastic driving inputs (see below). With regard to construct validity, we apply rDCM to empirical fMRI data and compare the results to those obtained by spectral DCM (Friston et al., 2014a), an alternative and established model of effective connectivity for rs-fMRI data.

### 2.2 Synthetic data

First, we evaluated the face validity of rDCM in simulation studies by generating synthetic rs-fMRI data for which the ground truth (i.e., the data-generating parameter values) were known. In particular, we assessed model parameter recovery in four different linear DCMs and evaluated performance as a function of the repetition time (TR) and signal-to-noise ratio (SNR) of the synthetic data. Each of the models comprised four regions, yet differed with regard to the degree of sparsity of the network architecture (**Fig. 1A**). More precisely, model 1 represented a full network where all regions were connected via reciprocal connections, model 2 comprised 75% of all possible connections, model 3 comprised 50% of all possible connections, and model 4 was the sparsest network with only 25% of all connections present. Please note that this simulation work was restricted to a small network with only four regions in order to allow comparability of rDCM with spectral DCM (Friston et al., 2014a), which represents an alternative variant of DCM for rs-fMRI data (see below).

**Figure 1.**
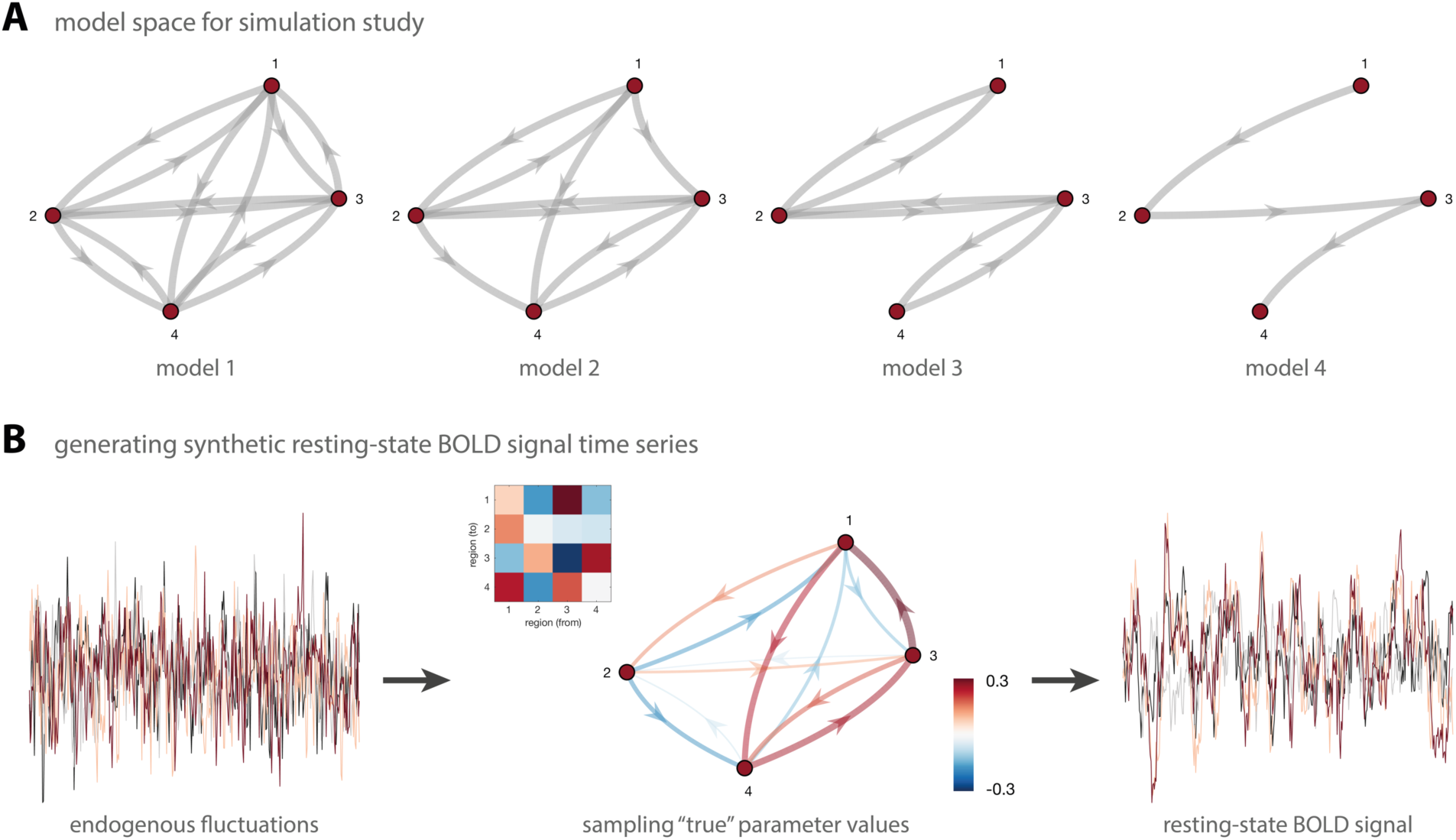
Pipeline for simulation studies on regression DCM (rDCM). **(A)** Model space for the simulation studies dedicated to assessing the face validity of rDCM for resting-state data. Four different network structures were utilized that varied in their degree of sparsity. Model 1 represents a fully connected model where all regions are coupled via reciprocal connections (i.e., 12 connectivity parameters), model 2 represents a network with 75% of all connections present (i.e., 9 connectivity parameters), model 3 represents a network with 50% of all connections present (i.e., 6 connectivity parameters), and model 4 represents the sparsest network with only 25% of all connectivity present (i.e., 3 connectivity parameters). **(B)** Regional endogenous fluctuations were generated from an AR(1) process and then utilized as driving inputs in the respective model. Specifically, we here used a classical deterministic DCM to generate synthetic data. Model parameters were sampled from their prior distributions and then utilized to generated synthetic BOLD signal time series that resembled the characteristic amplitudes and slow fluctuations observed during the resting state. Measurement noise, generated from another AR(1) process, was added on top of the predicted BOLD signal time series.

For each model, synthetic fMRI data were then generated to resemble the characteristic amplitude and low-frequency fluctuations seen in resting-state BOLD signal time series. To this end, we followed the procedure described in previous work (Friston et al., 2014a; Razi et al., 2015) and implemented in the MATLAB function *DEM_demo_induced_fMRI.m* as part of SPM12 (version R7487, Wellcome Centre for Human Neuroimaging, London, UK, http://www.fil.ion.ucl.ac.uk). In brief, endogenous neuronal fluctuations were generated independently for each region by using an AR(1) process with an autoregression coefficient of one half. The endogenous neuronal fluctuations were scaled to a standard deviation of 1/4. These values were identified by Friston et al. (2014a) to produce realistic BOLD signal changes of about 2%. Region-wise endogenous fluctuations were then utilized as driving inputs to a classical deterministic DCM (Friston et al., 2003). Similarly, measurement noise was generated independently for each region from another AR(1) process and added on top of the noise-free BOLD signal time courses. Synthetic resting-state fMRI data were simulated to mimic an experiment of 10 minutes duration. Importantly, BOLD signal time series were generated under a variety of different signal-to-noise ratios (SNR = [0.5 1 3]) and repetition times (TR = [2s, 1s, 0.5s]) in order to evaluate the performance of rDCM as a function of data quality and sampling rate, respectively. Here, SNR was defined as the ratio between the standard deviation of the signal and the standard deviation of the AR(1)-type measurement noise (i.e., *SNR* = *σ*_*signal*_/*σ*_*noise*_).

For each of the four models and each combination of SNR and TR, 20 different sets of observations (“synthetic subjects”) were generated. To this end, the generating (“true”) parameter values of each simulation were sampled from the prior distributions of the endogenous parameters. Parameter values too close to zero (i.e., |*a*_*i,j*_ | < 0.05) were discarded and re-sampled in order to ensure sufficient information transfer between regions.

Examples of endogenous fluctuations and ensuing synthetic BOLD signal time series are shown in **Fig. 1B**. Note that, as a result of the low-pass filtering of the hemodynamic response function, the BOLD signal is much smoother than the underlying endogenous fluctuations. The resulting BOLD signal time series are biologically realistic and show the characteristic slow (low-frequency) fluctuations typically observed during the resting state.

Based on the synthetic data, model inversion was performed by making use of the function *tapas_rdcm_estimate.m* from the rDCM toolbox as implemented in the **T**ranslational **A**lgorithms for **P**sychiatry **A**dvancing **S**cience (TAPAS) toolbox (www.translationalneuromodeling.org/software). Model parameter recovery was then assessed by computing (i) the root-mean-squared-error (RMSE) and (ii) the Pearson’s correlation coefficient (r) between the inferred and the generating (“true”) parameter values in order to quantify how much the estimated posterior parameter values differed from the ground truth.

### 2.3 Empirical data

Second, we examined the construct validity of rDCM in relation to spectral DCM. For this, we applied both models to an empirical rs-fMRI dataset and compared the results. Specifically, we made use of a large dataset collected by the Bipolar-Schizophrenia Network on Intermediate Phenotypes (B-SNIP-1) consortium, with one measure being rs-fMRI (Tamminga et al., 2013; 2014). Overall, participants with psychosis (including schizophrenia, schizoaffective disorder and psychotic bipolar I disorder), their first-degree relatives, and demographically matched healthy participants were collected for the B-SNIP-1 dataset.

#### 2.3.1 Participants

The data utilized in the present study were acquired by the B-SNIP-1 consortium as part of a large cross-sectional study of intermediate phenotypes in psychosis (Tamminga et al., 2013). The B-SNIP-1 consortium included five sites in the United States: (i) Baltimore, (ii) Chicago, (iii) Dallas, (iv) Detroit and Boston, and (v) Hartford. The sites used identical study protocols, and considerable effort was taken to harmonize recoding and testing conditions across sites by utilizing identical stimulus presentation and recording equipment. Furthermore, experimenters across sites were cross-trained and frequently monitored to ensure comparable data collection procedures. A detailed study description is provided elsewhere (Tamminga et al., 2013; 2014).

Given the methodological focus of this work, here we focus only on the healthy control participants (a comprehensive analysis of the full clinical sample will be presented in future work). Healthy participants were recruited from the local communities. They were without lifetime psychotic disorders and had no first-degree relatives with a history of psychotic or bipolar disorder according to the Family History Research Diagnostic Criteria (Andreasen et al., 1977). Overall, 459 healthy participants were assessed by the B-SNIP-1 consortium. For the current study, only those participants were included that had: (i) a complete data set comprising all relevant demographical information and neuroimaging data, and (ii) sufficiently high data quality of the fMRI measurements (exclusion criteria related to data quality are listed below). This yielded a final sample of 196 healthy participants that were included in our study (80 females, 116 males; age: 38.3±12.5 years; age range: 15-64 years). The study protocol was approved by the institutional review board at each local site. All participants provided written informed consent prior to inclusion and after they obtained a complete description of the study.

#### 2.3.2 Experimental procedure

While in the MR scanner, participants were asked to fixate on a small cross presented on the screen, and to remain alert with eyes open and head still. These instructions were designed to help prevent participants from falling asleep, provide an experimental control over visual inputs and eye movements, and reduce head movements. Head movements were further restricted by having participants’ heads placed in a custom-build head-coil cushion.

#### 2.3.3 Data acquisition

Structural and functional MRI data were acquired at the five sites of the B-SNIP-1 consortium. These included the University of Maryland School of Medicine (Baltimore), Commonwealth Research Center at Harvard Medical School (Boston), University of Illinois Medical Center (Chicago), University of Texas Southwestern Medical Center (Dallas), and Olin Neuropsychiatry Research Center at the Institute of Living (Hartford). Participants were scanned on 3 Tesla MR scanners of different manufacturers, including GE Signa, Siemens Trio, Philips Achieva, and Siemens Allegra. For comprehensive details on data acquisition parameters across the different sites, see Supplementary Material S1.

#### 2.3.4 Data preprocessing

Preprocessing of the MR data was performed using fMRIPrep (version 1.4.1; Esteban et al., 2019) which is based on Nipype (version 1.2.0; Gorgolewski et al., 2011) and was specifically designed for automated high-quality preprocessing in the context of large-scale datasets. We here utilized the standard preprocessing pipeline from fMRIPrep which comprised the following steps.

The T1-weighted (T1w) anatomical image was corrected for intensity non-uniformity (INU) with *N4BiasFieldCorrection* (Tustison et al., 2010), distributed with ANTs 2.2.0 (Avants et al., 2008), and used as T1w reference throughout the workflow. The T1w reference was then skull-stripped with a Nipype implementation of the *antsBrainExtraction* workflow. Brain tissue segmentation of cerebrospinal fluid (CSF), white matter (WM) and gray matter (GM) was performed on the brain-extracted T1w image using *fast* (Zhang et al., 2001), distributed with FSL 5.0.9 (Jenkinson et al., 2012). Volume-based spatial normalization to the MNI standard space was performed through nonlinear registration with *antsRegistration* (ANTs 2.2.0), using brain-extracted versions of both T1w reference and T1w template (MNI152NLin2009cAsym; Fonov et al., 2009).

Functional images were preprocessed by first generating a reference volume (BOLD reference) and its skull-stripped version by making use of a custom methodology of fMRIPrep. Head-motion parameters with respect to the BOLD reference (transformation matrices, and six corresponding rotation and translation parameters) were estimated before any spatiotemporal filtering using *mcflirt* (Jenkinson et al., 2002). Slice-timing correction was performed to account for differences in acquisition timing across slices. A deformation field to correct for susceptibility distortions was estimated based on the *fieldmap-less* approach in fMRIPrep (Wang et al., 2017). This procedure utilizes the T1w reference from the same subject as the undistorted target in a nonlinear registration scheme. To maximize the similarity between T2* contrast of the EPI scan and the T1w reference, the intensities of the latter are inverted. To regularize the optimization of the deformation field, displacements are constrained to be non-zero only along the phase-encoding direction and the magnitude of displacements is modulated with an average fieldmap template (Treiber et al., 2016). Based on the estimated susceptibility distortion, unwarped BOLD images were calculated for a more accurate co-registration with the anatomical reference by resampling functional images onto their original, native space through a single, composite transform to correct for head-motion and susceptibility distortions. This was followed by co-registration to the T1w reference using *flirt* (Jenkinson and Smith, 2001) with the boundary-based registration cost function (Greve and Fischl, 2009), implemented in FSL 5.0.9 (Jenkinson et al., 2012). Co-registration was configured with nine degrees of freedom to account for remaining distortions in the BOLD images. The corrected and coregistered functional images were then normalized into MNI standard space (MNI152NLin2009cAsym; Fonov et al., 2009) by making use of the warps obtained from the spatial normalization of the T1w reference (see above).

#### 2.3.5 Estimation of confound regressors

From the functional images, fMRIPrep computes several time series of potentially confounding influences that can be used subsequently (during the extraction of BOLD signal time series) as confound regressors to correct for variance of no interest. Here, we will focus only on those confound regressors that were utilized in the present analysis; for a comprehensive list of all confound regressors estimated by fMRIPrep, see elsewhere (Esteban et al., 2019).

In brief, framewise displacement (FD) and DVARS (Power et al., 2012) were calculated by making use of their implementations in Nipype, following the definitions by Power et al. (2014). Furthermore, three global signals were extracted within the CSF, WM, and a whole-brain mask. Additionally, the head motion estimates (translation and rotation) obtained during preprocessing were utilized as confound regressors. The confound regressors derived from head motion estimates and global signals were expanded with the inclusion of temporal derivatives and quadratic terms for each (Friston et al., 1996; Satterthwaite et al., 2013). Furthermore, frames that exceeded a threshold of 0.5mm for FD or 1.5 for standardized DVARS were annotated as motion outliers and marked using a stick regressor. Finally, regressors encoding a discrete cosine set with a 128s cut-off were utilized for high-pass filtering.

#### 2.3.6 Exclusion criteria

In addition to fMRIPrep, we utilized MRIQC (version 0.15.1; Esteban et al., 2017) for quality control of the neuroimaging data. Specifically, we ran MRIQC with the default parameters, with the exception of the framewise displacement threshold being adjusted to 0.5mm. Participants were excluded from further effective connectivity analysis if any of the following image quality metrics derived by MRIQC were above/below the specified threshold. For the structural images, exclusion criteria were defined as: (i) “*QI*_1_” > 0.005, (ii) “overlap_tpm_csf” < 0.1, (iii) “overlap_tpm_gm” < 0.3, and (iv) “overlap_tpm_wm” < 0.45. In brief, “*QI*_1_” represents the proportion of artifact-corrupted voxels as assessed using the algorithm described by Mortamet et al. (2009). Furthermore, “overlap_tpm_*” represents the overlap between the tissue probability maps (TPMs) estimated from the anatomical image and the corresponding maps from the MNI152NLin2009cAsym template. For the functional images, exclusion criteria were specified as: (i) “aqi” > 0.025, and (ii) “fd_perc” > 20.0. In brief, “aqi” is AFNI’s mean quality index computed by the *3dTqual* routine, and “fd_perc” represents the percentage of scans with a FD above the FD threshold with respect to the entire time series. A full description of the metrics is available from https://mriqc.readthedocs.io/en/latest/measures.html.

#### 2.3.7 Time series extraction: small networks

To evaluate the construct validity of rDCM in application to resting-state fMRI data, we examined functional integration during the resting state from two perspectives: modes and nodes in rs-fMRI (Friston et al., 2014b). First, we examined effective connectivity among key intrinsic networks of the human brain (i.e., modes). Specifically, we investigated the established anticorrelations between the task-negative default mode network, engaged during rest and internally directed tasks, and task-positive networks, engaged during externally oriented tasks. These anticorrelations have been proposed to serve as a fundamental characteristic of endogenous or spontaneous fluctuations during the resting state (Fox et al., 2005; Raichle, 2015; Smith et al., 2009; but see Discussion and Fox et al., 2009).

To this end, we selected in a first step four modes of interest, comprising the default mode network (DMN; Buckner et al., 2008; Raichle, 2015), dorsal attention network (DAN; Corbetta and Shulman, 2002; Fox et al., 2006), salience network (SAN; Dosenbach et al., 2007; Menon and Uddin, 2010), and central executive network (CEN; Spreng et al., 2013; 2010). Masks for each of the four RSNs were created by making use of previously published templates (Shirer et al., 2012) and are displayed in **Fig. 2A**.

**Figure 2.**
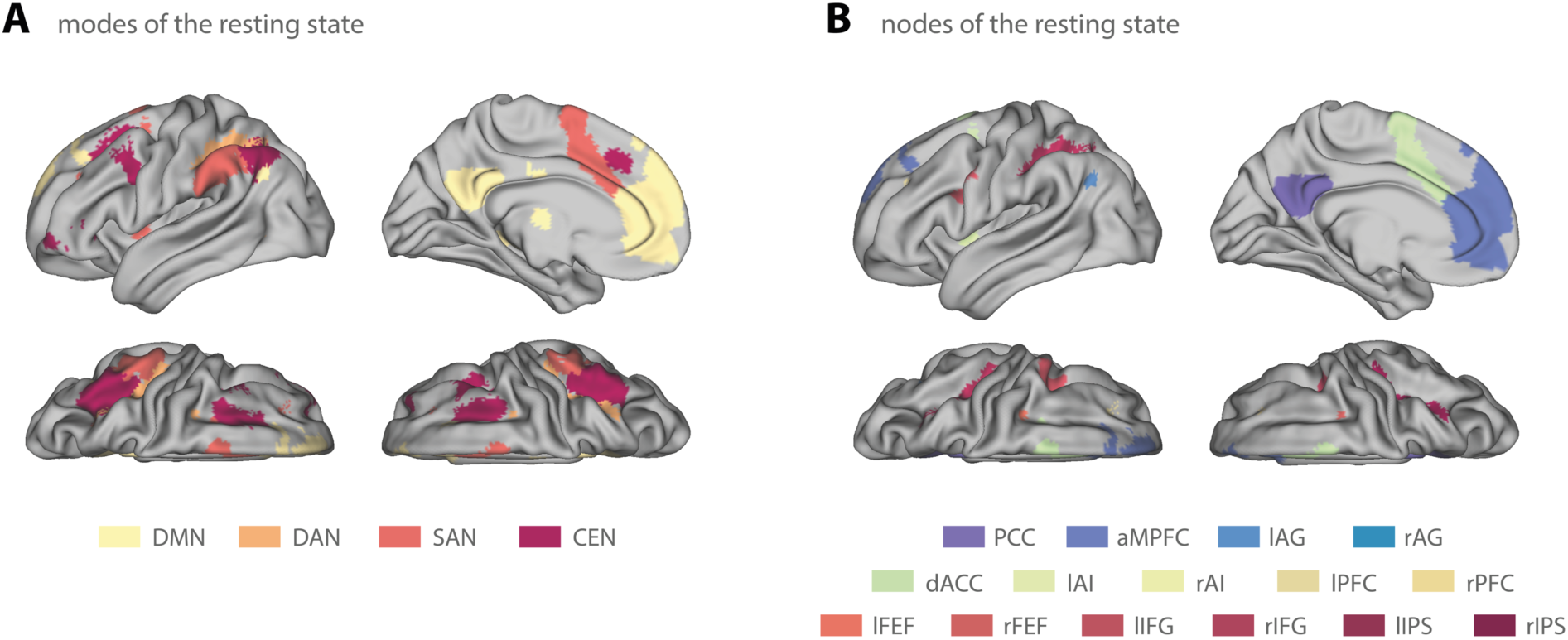
Masks for the empirical analysis of the B-SNIP-1 resting-state data. **(A)** Masks comprising four key intrinsic networks (i.e., modes) of the resting state. Specifically, we included the default mode network (DMN; *yellow*), dorsal attention network (DAN; *orange*), salience network (SAN; *red*), and the central executive network (CEN; *purple*). **(B)** Masks comprising 15 subcomponents of three key intrinsic networks (DMN, DAN, and SAN) of the resting state. For the DMN (*blueish colors*): posterior cingulate cortex (PCC), anterior medial prefrontal cortex (aMPFC), and left and right angular gyrus (lAG and rAG). For the SAN (*yellowish colors*): dorsal anterior cingulate cortex (dACC), left and right anterior insula (lAI and rAI), and left and right anterior prefrontal cortex (lPFC and rPFC). For the DAN (*reddish colors*): left and right frontal eye fields (lFEF and rFEF), left and right inferior frontal gyrus (lIFG and rIFG), and left and right superior parietal lobe (lSPL and rSPL). All masks were taken from the templates published in Shirer et al. (2012).

In a second step, we investigated effective connectivity during the resting state not only at the level of entire networks, but at the level of the regional components (nodes) of those networks. To this end, we restricted ourselves to the DMN, DAN, and SAN, in line with previous work on the hierarchical organization of RSNs by Zhou et al. (2018). Consistent with Zhou et al. (2018), we subdivided the three key intrinsic networks as follows: For the (core) DMN, we identified 4 regions: posterior cingulate cortex (PCC), anterior medial prefrontal cortex (aMPFC), and left and right angular gyrus (lAG and rAG). For the DAN, we identified 6 regions: left and right frontal eye fields (lFEF and rFEF), left and right inferior frontal gyrus (lIFG and rIFG), and left and right superior parietal lobe (lSPL and rSPL). For the SAN, we identified 5 regions: dorsal anterior cingulate cortex (dACC), left and right anterior insula (lAI and rAI), and left and right anterior prefrontal cortex (lPFC and rPFC). This yielded a total of 15 regions of interest (ROIs) from which BOLD signal time series were extracted. Again, we utilized the templates provided by Shirer et al. (2012) as masks for the respective ROIs (**Fig. 2B**).

Delineating the components of the intrinsic networks allowed for a more fine-grained analysis of effective connectivity during the resting state and for disentangling functional integration within *and* between networks. It also served as a more challenging test scenario for rDCM given the increase in network size and, thus, number of free connectivity parameters. Having said this, the resulting network sizes are still an order of magnitude smaller than what is possible using rDCM. However, for the assessment of construct validity, we deliberately restricted ourselves to relatively small networks in order to allow for a comparison between rDCM and spectral DCM (but see below for an application of rDCM to rs-fMRI data from a whole-brain network with more than 200 regions).

For either of the two analysis pipelines (i.e., “modes” and “nodes”), BOLD signal time series were extracted as the first eigenvariate of all voxels within the respective mask. Time series were mean centered and confound-related variance was removed (by regression using all nuisance regressors included in the first-level GLM).

#### 2.3.8 Time series extraction: whole-brain network

Finally, to demonstrate that rDCM can infer effective connectivity patterns during the resting state at the whole-brain level, we utilized the Human Brainnetome atlas as a whole-brain parcellation scheme (Fan et al., 2016). The Brainnetome atlas represents a connectivity-based parcellation derived from diffusion weighted imaging (DWI) data. In brief, the atlas comprises 246 distinct parcels (123 per hemisphere), including 210 cortical and 36 subcortical regions. Due to signal dropouts in the raw functional images (especially in inferior temporal regions near the skull base), BOLD signal time series could not be extracted for all regions defined by the Brainnetome atlas. Overall, 221 regions could be extracted in all participants. However, in order to ensure interhemispheric consistency of the network, we further reduced this set by excluded regions that were present in only one hemisphere.

This yielded a total of 212 brain regions from which sensible BOLD signal time series could be obtained in every participant. As for the small-network analyses, BOLD signal time series were then extracted as the first eigenvariate of all voxels within the respective mask. Time series were mean centered and confound-related variance was removed (by regression using all nuisance regressors included in the first-level GLM).

#### 2.3.9 rDCM analysis

The extracted BOLD signal time series from the previous step were then utilized for subsequent effective connectivity analyses using rDCM. For this, we constructed fully connected networks where all modes/regions were coupled to each other via reciprocal connections. This yielded a total of 16 free parameters for the 4-mode network and 225 free parameters for the 15-node network (all possible inter-regional connections and inhibitory self-connections), as well as 18,260 free parameters for the whole-brain network (all inter-regional connections present in the structural connectome of the Brainnetome atlas and inhibitory self-connections) to be estimated. Again, model inversion was performed by utilizing the routine *tapas_rdcm_estimate.m* from the rDCM toolbox as implemented in TAPAS.

## 3 Results

### 3.1 Simulations

#### 3.1.1 Face validity

First, we performed comprehensive simulation studies to assess the face validity of rDCM for modeling resting-state fMRI data. To this end, we tested whether rDCM can recover generating (“true”) parameter values in four different 4-region networks (**Fig. 1**) under various settings of SNR and TR of the synthetic data.

Overall, model parameter values could be recovered faithfully over a wide range of SNR and TR settings, for all four network architectures (**Fig. 3A** and **Supplementary Fig. 1**). Specifically, root-mean-squared-errors (RMSE) were below 0.15 in all tested cases. On average, we found the mean RMSE to range from 0.09 (95% confidence interval: [0.08 0.10]) for model 1 to 0.04 [0.037 0.045] for model 4 for the most challenging case of high measurement noise (SNR = 0.5) and low sampling rate (TR = 2s). Note that the absolute RMSE value is difficult to interpret as it will depend on the scaling of the data; hence, values should only be interpreted in a relative manner as they allow comparison across different data settings (i.e., SNR, TR) as well as across the different DCM variants. The average Pearson’s correlation coefficient 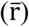 ranged from 0.70 [0.63 0.76] for model 1 to 0.95 [0.92 0.97] for model 4 for the low-SNR-slow-TR scenario, and thus showed a pattern highly consistent with the RMSE. Notably, averaging was performed by (i) transforming individual correlation coefficients to z-space using Fisher r-to-z transformation, (ii) computing mean as well as lower and upper bound of the 95% confidence interval in z-space, and finally (iii) back-transforming estimates to r-space. Overall, we observed an (expected) dependence of model parameter recovery on the sparsity of the network, with the most accurate results being observed for the simplest model 4 with the lowest number of free parameters (**Fig. 3A**). Furthermore, we observed a strong dependence of model parameter recovery performance on the data quality, suggesting that rDCM can infer upon connection strengths more faithfully for higher SNR settings of the (synthetic) BOLD signal time series. A similar effect was observed for the sampling rate (i.e., TR). Specifically, model parameter recovery performance of rDCM improved for faster TR settings, although this was somewhat less pronounced as the effect of the SNR (see **Supplementary Fig. 1**). The observed dependencies of model parameter recovery performance on SNR and TR settings are consistent with previous simulation analyses on rDCM in the context of task-based data (Frässle et al., 2018a; 2017).

**Figure 3.**
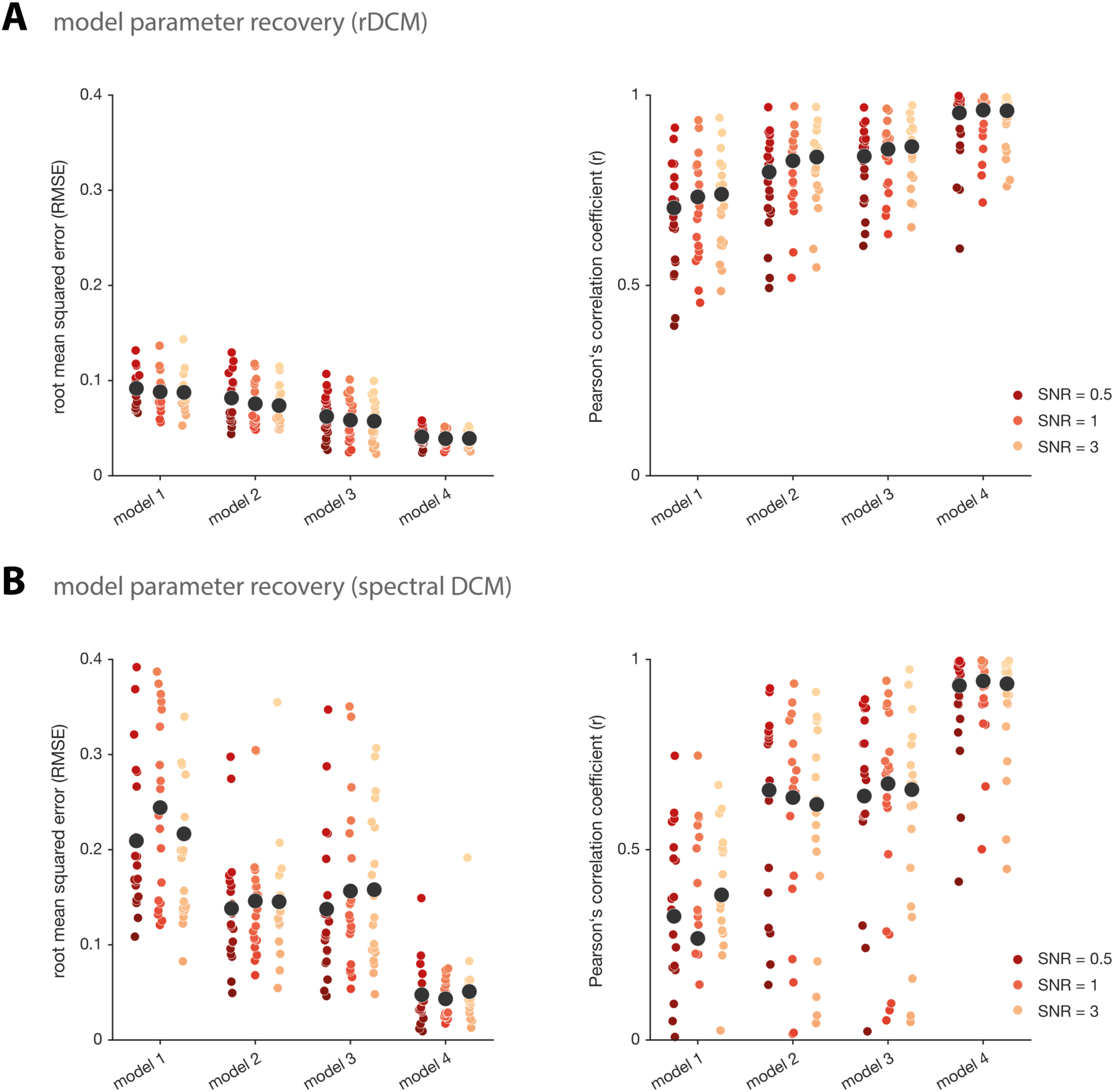
Model parameter recovery for regression DCM and spectral DCM. **(A)** Model parameter recovery for synthetic resting-state fMRI data in terms of the root mean squared error (*left*) and the Pearson’s correlation coefficient (*right*) between the inferred and the “true” data-generating parameter values for regression DCM (rDCM), and **(B)** spectral DCM. Each colored dot represents the respective RMSE or Pearson’s correlation coefficient for an individual synthetic subject. Black dots represent the mean RMSE or mean Pearson’s correlation coefficient, averaged across all 20 synthetic subjects. Note that averaging of the individual Pearson’s correlation coefficients was performed in z-space (see main text). Results are shown for four different network architectures, where model 1 is the most complex (full) model and model 4 is the sparsest network. Furthermore, results are shown for an exemplary repetition time (TR) of 2s and three different signal-to-noise ratios (SNRs): 0.5 (red), 1 (orange), and 3 (yellow).

#### 3.1.2 Comparison to spectral DCM

In a second step, we compared the model parameter recovery performance of rDCM with results obtained with spectral DCM (Friston et al., 2014a), which represents an alternative variant of DCM for fMRI that is suited to model resting state data. Importantly, spectral DCM has already been assessed in terms of face validity (Friston et al., 2014a), construct validity (Razi et al., 2015), and test-retest reliability (Almgren et al., 2018). Consequently, spectral DCM serves as a useful benchmark against which rDCM can be challenged.

Spectral DCMs were fitted to the exact same synthetic BOLD signal time series that have previously been utilized for rDCM. As before, model parameter recovery of spectral DCM improved as a function of network sparsity (**Fig. 3B**). However, the dependence of model parameter recovery on the characteristics of the synthetic fMRI data was less clear for spectral DCM: while no obvious dependence on data quality (i.e., SNR) could be observed, model parameter recovery showed a tendency to improve as a function of sampling rate (see **Supplementary Fig. 2**), which is consistent with previous work (Friston et al., 2014a). When comparing parameter recovery performance across the two DCM variants, overall, rDCM performed better than spectral DCM. More specifically, rDCM and spectral DCM were on par for the simplest model 4, whereas for all other network architectures, rDCM outperformed spectral DCM in terms of RMSE and Pearson’s correlation coefficient. This pattern was observed across all SNR and TR settings.

In summary, our simulation analyses provide evidence for the face validity of rDCM in application to synthetic resting-state fMRI data in a relatively broad range of SNR and TR settings (**Supplementary Fig. 1-2**).

### 3.2 Empirical analyses: 4-mode network

#### 3.2.1 Effective connectivity during resting state

Next, we investigated the construct validity of rDCM in comparison to spectral DCM, using an empirical rs-fMRI dataset. We here used the 196 healthy participants from the B-SNIP-1 consortium (Tamminga et al., 2013) described earlier (see Methods and Materials).

First, we investigated effective connectivity among four key modes of the resting state (**Fig. 2A**), namely the default mode network (DMN), dorsal attention network (DAN), salience network (SAN), and central executive network (CEN). Individual connectivity parameters were estimated in a fully connected network using rDCM. Model inversion resulted in biologically plausible connectivity patterns (**Fig. 4A**). Specifically, we observed pronounced inhibitory afferent (inward) and efferent (outward) connections between the DMN, representing the task-negative network, and the DAN and SAN, representing task-positive networks. Additionally, we observed excitatory reciprocal connections between DAN and SAN. Interestingly, the CEN exerted a relatively strong excitatory influence on the DMN and simultaneously received a (weaker) positive influence from the DMN. Furthermore, the CEN was coupled positively to the DAN and negatively to the SAN.

**Figure 4.**
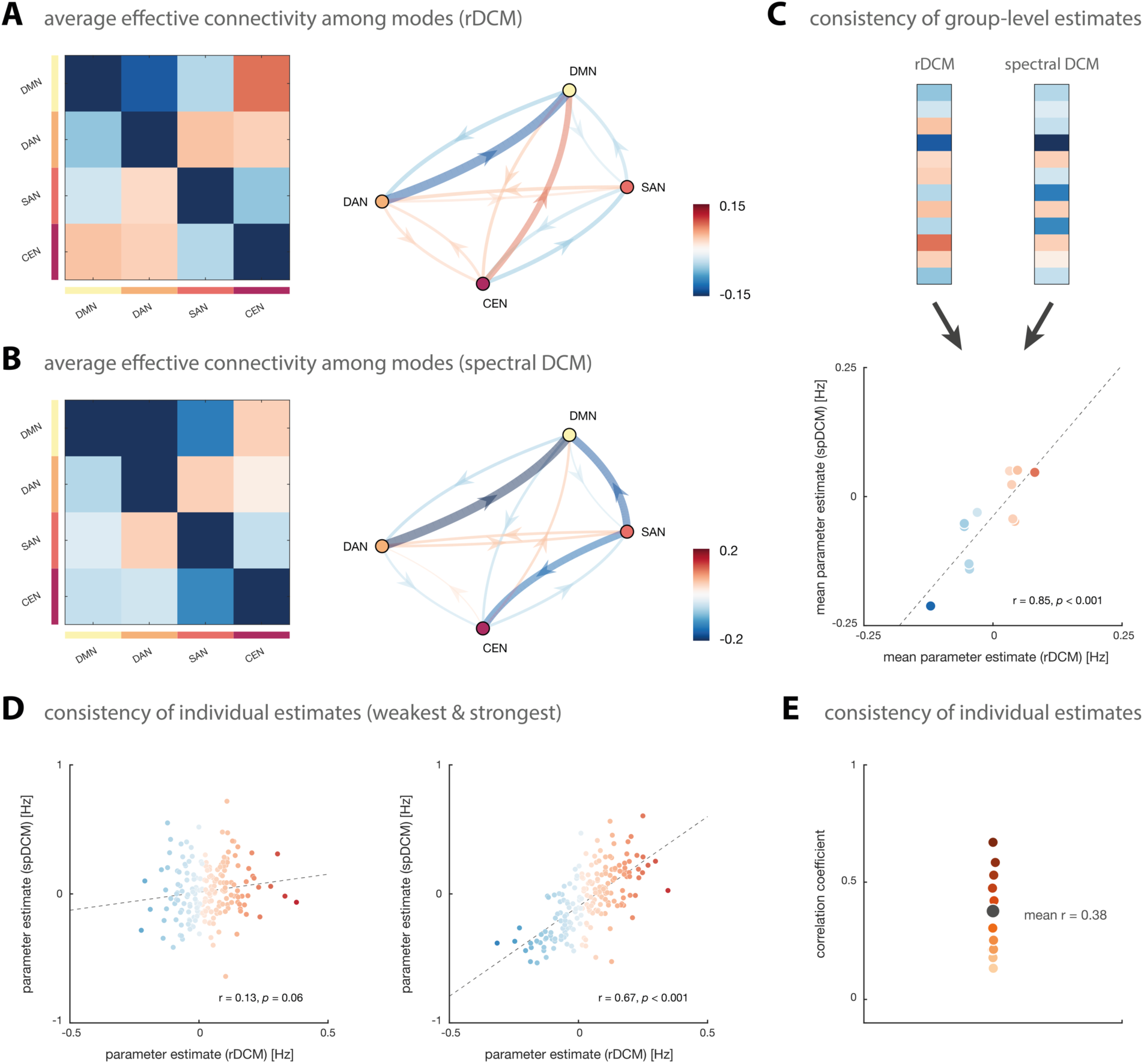
Effective connectivity amongst key modes of the resting state. **(A)** Group-averaged posterior parameter estimates during the resting state as inferred with regression DCM (rDCM), and **(B)** spectral DCM (spDCM). Note that in order to make the correlation between group-averaged connectivity estimates of the two DCM variants more easily visible, color was scaled in relation to the maximal values of each method. Hence, we utilize slightly different color scales for the two DCM variants for illustration purposes. **(C)** Consistency of group-level parameter averages between rDCM and spDCM. Shown are the group-averaged inter-regional connectivity estimates (*top*) as well as the Pearson’s correlation coefficient (r) amongst them (*bottom*). Dots are colored using the rDCM color scale. **(D)** Consistency of individual parameter estimates for each connection separately; here, shown for the connection with the weakest (*left*) and strongest association (*right*) between the two DCM variants. Dots are colored using the rDCM color scale. **(E)** Range of correlation coefficients across all 12 connections of the 4-mode network. Each colored dot represents the Pearson’s correlation coefficient for an individual inter-regional connection and the black dot denotes the mean Pearson’s correlation coefficient, averaged across all connections. Note that averaging of individual Pearson’s correlation coefficients was performed in z-space (see main text).

Based on the posterior parameter estimates, one can compute a metric of hierarchy strength as the difference between averaged absolute efferent and afferent connections for each of the four intrinsic networks of the resting state. This approach has been used by a previous DCM study on the hierarchical organization of resting-state networks (Zhou et al., 2018), and is similar to approaches used for analyzing hierarchical projections in the monkey brain (Goulas et al., 2014) and the hierarchical organization of the prefrontal cortex in humans (Nee and D’Esposito, 2016). Importantly, this definition of “hierarchy” should not be confused with other accounts that relate to layer-specificity of forward and backward connections (Felleman and Van Essen, 1991; Markov et al., 2014b). Instead, the present definition simply refers to the degree of influence that one mode exerts over others (for a comprehensive review of alternative definitions of hierarchy, see Hilgetag and Goulas, 2020). The total hierarchy strength was indicative of a hierarchical organization of the four networks with the DMN at the bottom (−0.12), predominantly serving as a sink, and the other task-positive networks ranking higher (DAN: 0.05, SAN: 0.02, CEN: 0.05), predominantly serving as neuronal drivers. This finding is consistent with previous observations on the hierarchical organization of resting-state networks (Sridharan et al., 2008; Zhou et al., 2018).

#### 3.2.2 Comparison to spectral DCM

As for the simulations above, we again compared our findings from rDCM to those obtained by spectral DCM, using the identical BOLD signal time series. We observed that, at the group level, effective connectivity patterns were highly consistent across the two DCM variants (**Fig. 4B**). In brief, spectral DCM also revealed pronounced inhibitory afferent and efferent connections between the DMN and the DAN and SAN, as well as excitatory reciprocal connections amongst DAN and SAN. The only (qualitative) difference between spectral DCM and rDCM results was observed in how the CEN integrated into this network. While, consistent with the rDCM results, efferent connections from the CEN to the DMN and DAN were excitatory, afferent connections to the CEN from DMN and DAN were inhibitory. With regard to the coupling between CEN and SAN, spectral DCM and rDCM again yielded consistent results. This similarity between the two DCM variants was also represented by a very high Pearson’s correlation coefficient between the group-level inter-regional connectivity estimates (r = 0.85, *p* < 0.001; **Fig. 4C**).

In a second step, we investigated the consistency of parameter estimates at the individual level (rather than at the group level) for each connection separately. At the individual level, connections showed some variability in the degree of consistency between the two DCM variants. While for some connections, the Pearson’s correlation between rDCM and spectral DCM parameter estimates was relatively weak, as seen for the connection from CEN to DAN (r = 0.13, *p* = 0.06; **Fig. 4D**, *left*), others showed much higher consistency, like the connection from DAN to CEN (r = 0.67, *p* < 0.001; **Fig. 4D**, *right*). Overall, the mean Pearson’s correlation coefficient, averaged across connections, was 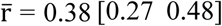 (**Fig. 4E**). Again, averaging was performed by (i) transforming individual correlation coefficients to z-space using Fisher r-to-z transformation, (ii) computing mean as well as lower and upper bound of the 95% confidence interval in z-space, and finally (iii) back-transforming estimates to r-space. Despite the observed variability, our results suggest that, across connections, there is significant evidence for consistency across parameter estimates from the two DCM variants at the individual level, as assessed using a one-sample *t*-test after Fisher r-to-z transformation of the correlation coefficients (t_(1,11)_ = 6.40, *p* < 0.001).

### 3.3 Empirical analysis: 15-node network

#### 3.3.1 Effective connectivity during resting state

Next, we aimed to study effective connectivity during the resting state not only at the level of entire networks, but at the level of the regional components of those networks. To this end, we restricted our analysis to the DMN, DAN, and SAN and, consistent with Zhou et al. (2018), divided these three networks into 15 subcomponents (**Fig. 2B**). This yielded four regions for the DMN (PCC, aMPFC, lAG, and rAG), five regions for the SAN (dACC, lAI, rAI, lPFC, and rPFC), and six regions for the DAN (lFEF, rFEF, lIFG, rIFG, lSPL, and rSPL).

As for the 4-mode network, individual connectivity parameters were estimated in a fully connected network using rDCM, which resulted in biologically plausible connectivity patterns (**Fig. 5A**). The first observation that becomes apparent from our result is the clear modular structure of the average endogenous connectivity matrix where regions that belong to the same RSN group together and show almost exclusively positive (excitatory) within-network interactions. Conversely, between-network connections (i.e., connections originating from a region in one RSN and terminating at a region in another RSN) were overall weaker and more variable, showing both inhibitory and excitatory influences. More precisely, afferent and efferent connections between regions of the DMN and regions of the SAN and DAN were still predominantly negative, whereas connections between the two task-positive networks were more diverse.

**Figure 5.**
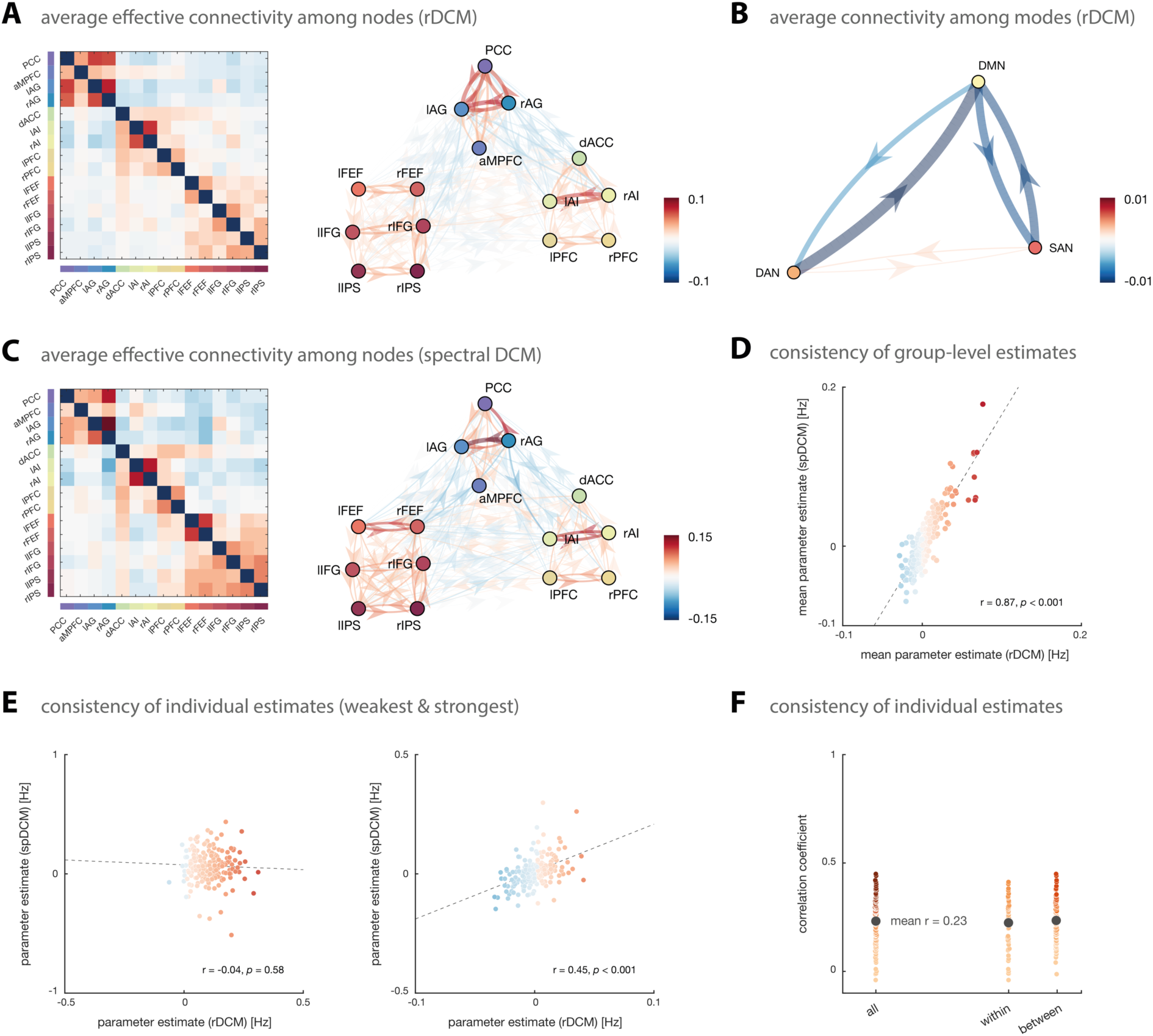
Effective connectivity amongst key brain regions of the resting state. **(A)** Group-averaged posterior parameter estimates during the resting state as inferred with regression DCM (rDCM). **(B)** Average connection strength between resting-state network (i.e., DMN, DAN, and SAN), computed as the average of the strength of all connections originating in one resting-state network and terminating in another one. **(C)** Group-averaged posterior parameter estimates during the resting state as inferred with spectral DCM (spDCM). Note that in order to make the correlation between group-averaged connectivity estimates of the two DCM variants more easily visible, color was scaled in relation to the maximal values of each method. Hence, we utilize slightly different color scales for the two DCM variants for illustration purposes. **(D)** Consistency of group-level parameter averages between rDCM and spDCM. Shown is the Pearson’s correlation coefficient (r) amongst the group-averaged inter-regional connectivity estimates. Dots are colored using the rDCM color scale. **(E)** Consistency of individual parameter estimates for each connection separately; here, exemplarily shown for the connection with the weakest (*left*) and strongest association (*right*) between the two DCM variants. Dots are colored using the rDCM color scale. **(F)** Range of correlation coefficients across all 210 connections of the 15-node network, as well as separately for all connections *within* a resting-state network and all connections *between* resting state network. Each colored dot represents the Pearson’s correlation coefficient for an individual inter-regional connection and the black dot denotes the mean Pearson’s correlation coefficient, averaged across all connections. Note that averaging of individual Pearson’s correlation coefficients was performed in z-space (see main text).

In order to allow for direct comparability between this analysis of nodes and the previous analysis of modes, we computed the average connection strength between networks (**Fig. 5B**). For example, the strength of all connections originating from a node of the DAN and terminating at a node of the DMN were averaged in order to yield the between-network connection strength from the DAN to the DMN. This analysis suggested that, at the between-network level, results from the 15-node network were highly consistent with the ones obtained in the previous analysis of the 4-mode network. Specifically, we again observed that the DMN was reciprocally connected to the other two RSNs via strong inhibitory connections, whereas the two task-positive networks (i.e., DAN and SAN) showed exhibitory coupling (although very weakly).

Finally, based on these between-network connections (**Fig. 5B**), we again computed the hierarchy strength as the difference between averaged absolute efferent and afferent connections between the three RSNs. As for the previous analysis, the total hierarchy strength was indicative of a hierarchical organization of the three networks with the DMN at the bottom (−0.007), predominantly serving as a sink, and the two task-positive networks ranking higher (DAN: 0.006, SAN: 0.0007), predominantly serving as neuronal drivers. In fact, the three RSNs displayed even the exact same ordering as in the previous analysis of modes – a remarkable observation given the differences in the underlying networks – and are thus again consistent with previous work (Sridharan et al., 2008; Zhou et al., 2018).

#### 3.3.2 Comparison to spectral DCM

As before, we compared our empirical findings from rDCM against the results obtained when utilizing spectral DCM to fit the identical BOLD signal time series. Group-level effective connectivity patterns were again highly consistent across the two DCM variants (**Fig. 5C**). Specifically, spectral DCM revealed the same modular structure of the (average) endogenous connectivity matrix with pronounced excitatory connections between regions of the same RSN and a more diverse pattern for between-network connections. The similarity between the two DCM variants was again represented by a very high Pearson’s correlation coefficient between the group-level connectivity estimates (r = 0.87, *p* < 0.001; **Fig. 5D**).

At the individual level, connections again varied in the degree of consistency between the two DCM variants. Correlations between rDCM and spectral DCM parameter estimates ranged from virtually absent (r = −0.04, *p* = 0.58) for the connection from left to right IFG (**Fig. 5E**, *left*) to robust (r = 0.45, *p* < 0.001) for the connection from left IPS to dACC (**Fig. 5E**, *right*), representing a significant correlation. Overall, the mean correlation coefficient, averaged across all connections, was 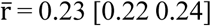 (**Fig. 5F**), and thus slightly decreased as compared to the 4-mode networks. Nevertheless, consistency between rDCM and spectral DCM was significantly larger than zero, as assessed using a one-sample *t*-test (t_(1,209)_ = 34.67, *p* < 0.001). Notably, consistency between the two DCM variants was not significantly different for within-module as compared to between-module connections (t_(1,208)_ = −0.75, *p* = 0.46), suggesting that there was no systematic difference between those two types. As before, averaging of correlation coefficients as well as statistical testing were performed in z-space – that is, after Fisher r-to-z transformation of the individual correlation coefficients.

In summary, these results were very similar to the ones obtained for the 4-mode networks, although correlations at the individual level were overall somewhat lower for the 15-node networks.

### 3.4 Empirical analysis: whole-brain network

Having established the face validity and construct validity of rDCM for rs-fMRI data in small networks, we assessed the practical utility of rDCM for inferring resting-state effective connectivity at the whole-brain level. To this end, we utilized the Brainnetome atlas as a whole-brain parcellation scheme, using a 212-node network covering the entire cortex (see Methods for details). Structural connectivity information provided by the Brainnetome atlas was used to constrain the network, resulting in 18,260 directed connections to be estimated (**Fig. 6A**).

**Figure 6.**
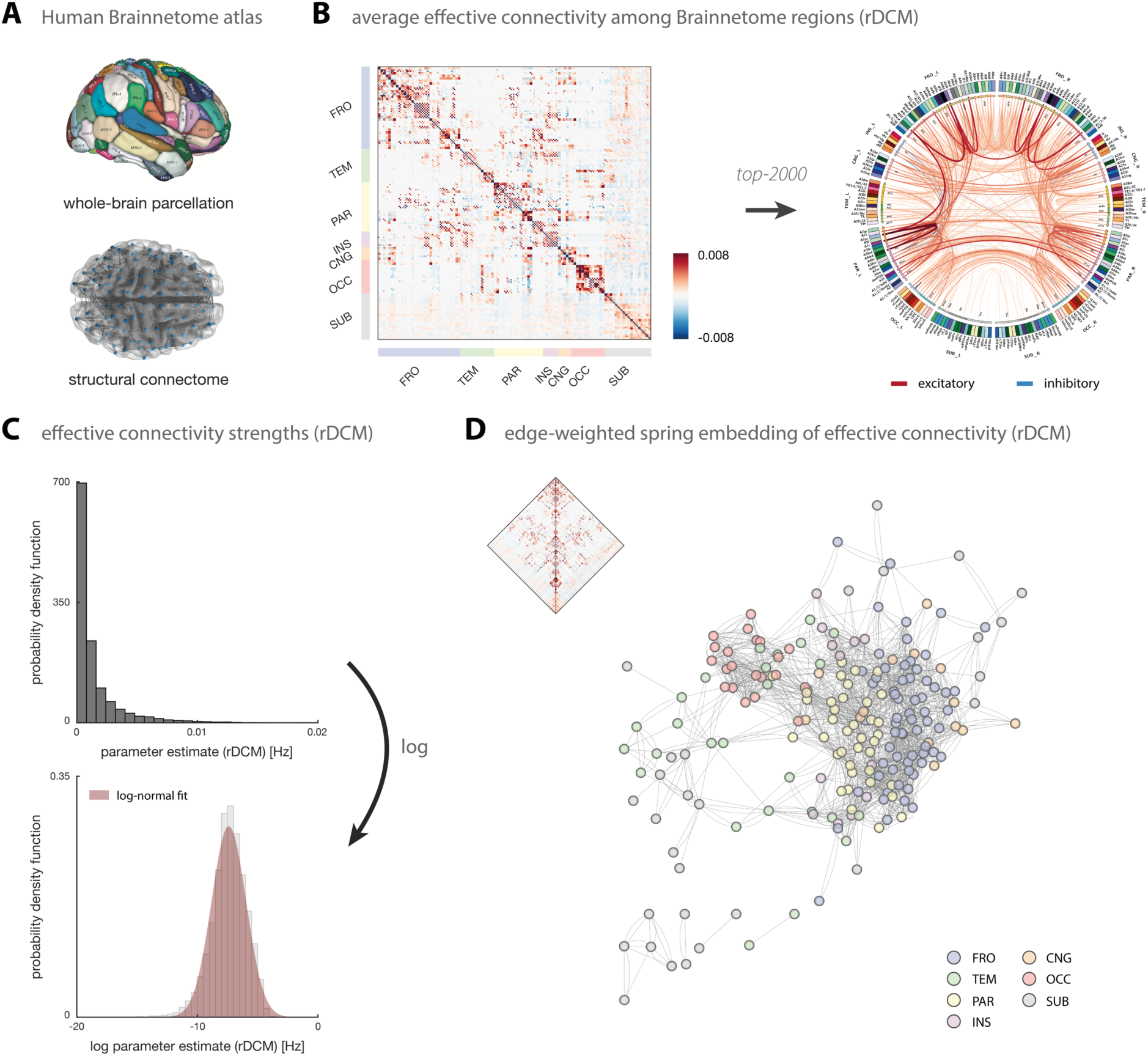
Whole-brain effective connectivity during the resting state. **(A)** The whole-brain rDCM analysis used the Human Brainnetome atlas that provides both a whole-brain parcellation of cortical and subcortical regions as well as information on the structural connectivity amongst these parcels. The image of the Brainnetome parcellation was reprinted with permission from Fan et al. (2016). **(B)** Group-averaged posterior estimates of directed connection strengths during the resting state as inferred with rDCM. Results are displayed as the full adjacency matrix (*left*) and as a connectogram of the top 2,000 connections with highest (absolute) weights (*right*). The connectogram was produced using Circos (publicly available at http://www.circos.ca/software/). Regions are divided into distinct sets, including frontal (FRO), temporal (TEM), parietal (PAR), insular (INS), cingulate (CNG)^1^, occipital (OCC), and subcortical (SUB). **(C)** Distribution of (absolute) endogenous connectivity parameters both in native (*top*) and logarithmic space (*bottom*). **(D)** Edge-weighted spring embedding of the endogenous connectivity matrix based on the top 2000 connections (see above). The edge-weighted spring embedding projection was computed using Cytoscape (publicly available at https://cytoscape.org/download.html).

Applying rDCM to rs-fMRI data from these 212 regions resulted in biologically plausible connectivity patterns (**Fig. 6B**). The average endogenous connectivity matrix displayed a clear modular structure where regions within the same lobe (e.g., frontal, occipital) were strongly connected amongst each other (**Fig. 6B**, *left*). Specifically, connections amongst regions within the same lobe were stronger and mostly positive (excitatory), whereas connections amongst regions in different lobes were more variable (i.e., showing both inhibitory and excitatory influences). This modular structure at the whole-brain level is consistent with our analysis of the connectivity amongst resting-state nodes. Furthermore, we found that while the majority of strong connections are within hemispheres, we also observed strong interhemispheric interactions, primarily amongst homotopic regions (**Fig. 6B**, *right*).

Inspection of the distribution of (absolute) endogenous connectivity parameters revealed a log-normal weight distribution, with connection strengths varying over several orders of magnitude (**Fig. 6C**). This suggests a predominance of weak connections (corresponding to low-frequency influences) in the average endogenous connectivity matrix. At the same time, the log-normal distribution highlights the presence of an extended tail, indicating many strong connections (i.e., higher-frequency influences) in the effective connectome. This finding is consistent with previous reports on log-normal distributions of anatomical connection weights in the mouse cortex (Oh et al., 2014; Wang et al., 2012b) and in the macaque cortical connectome (Markov et al., 2013; 2014a). The fall-off from lower to higher frequency influences in the distribution of rDCM parameter estimates is also broadly consistent with the general spectral characteristics of rs-fMRI data where low frequencies dominate (Chen and Glover, 2015; Kalcher et al., 2014; Niazy et al., 2011).

Finally, the modular structure of the resting-state effective connectivity becomes even more apparent when displaying the endogenous connectivity matrix as an edge-weighted spring-embedded projection. In brief, edge-weighted spring embedding positions nodes that interact strongly next to each other, by treating each node as a steel ring linked by springs (edges), thereby forming a mechanical system (Kamada and Kawai, 1989). The optimal layout is then defined as one that minimizes the total energy of the system. Applying edge-weighted spring embedding to the top 2000 connections of the average endogenous connectivity matrix revealed a clear modular structure with a tendency for nodes from the same set to cluster together (**Fig. 6D**). This was particularly clear for regions in the occipital (*red*), parietal (*yellow*), and frontal lobe (*blue*). Furthermore, the spring-embedded projection of the endogenous connectivity matrix was indicative of a hierarchical organization from occipital over parietal to frontal regions. This is consistent with the widespread view on brain organization as a hierarchy ranging from unimodal (e.g., visual) to transmodal (e.g., frontal) regions (Mesulam, 1998). Regions from some lobes (e.g., temporal) did not integrate clearly into this hierarchy, probably due to a dominance of weaker connection strengths below the chosen threshold. Notably, the spring-embedded projection shown here should only be treated as a qualitative illustration, as the exact layout will depend on, for instance, the density of the network (and thus thresholding) and the exact initialization of the edge-weighted spring embedding algorithm. Having said this, we verified that the overall pattern observed here holds for a range of different settings.

### 3.5 Computational burden

To provide an overview of computational efficiency, we report the run-time for the different analyses. These values should be treated as a rough indication only, as run-times will depend on the specific hardware used. Here, for all analyses, we used a single processor core on the Euler cluster at ETH Zurich (https://scicomp.ethz.ch/wiki/Euler).

First, rDCM took roughly 1 second for the small networks, regardless of the network architecture. More specifically, for the 4-mode networks (16 free parameters: 12 connection and 4 inhibitory self-connection parameters), model inversion took on average 0.48±0.07s (range: 0.35-0.77s), whereas for the 15-node networks (225 free parameters: 210 connection and 15 inhibitory self-connection parameters), model inversion took on average 0.62±0.10s (range: 0.47-1.32s). For the whole-brain networks (18,260 free parameters: 18,048 connection and 212 inhibitory self-connection parameters), model inversion took on average 132.30±20.26s (range: 90.63-175.30s). This highlights the computational efficiency of rDCM in application to rs-fMRI data and illustrates that the model scales gracefully with the number of regions, consistent with previous work on task-fMRI (Frässle et al., 2018a; 2017).

In comparison, while spectral DCM was also computationally efficient for the small networks, run-times were considerably longer. Specifically, for the 4-mode networks (30 free parameters: 12 connection, 4 inhibitory self-connection, 6 hemodynamic and 8 endogenous noise parameters), model inversion took on average 14.07±5.99s (range: 8.88-48.26s), whereas for the 15-node networks (261 free parameters: 210 connections, 15 inhibitory self-connections, 17 hemodynamic and 19 endogenous noise parameters), model inversion already took on average 3031±1551s (range: 819-12157s).

In summary, although the number of free parameters of the small networks was comparable for the two DCM variants, run-times were up to three orders of magnitude larger for spectral DCM. Specifically, our run-time analyses suggest that rDCM scales much more gracefully to large networks as compared to spectral DCM. Here, we only applied rDCM to rs-fMRI data from a parcellation with more than 200 areas; this was because runtimes of 21-42 hours per subject were reported for spectral DCM in application to a much smaller network of 36 regions (Razi et al., 2017).

## 4 Discussion

In this paper, we assessed the face validity and construct validity of regression DCM (rDCM; Frässle et al., 2018a; 2017) for inferring effective connectivity during the resting state (i.e., unconstrained cognition) from non-invasive fMRI data. We demonstrated in simulation studies that rDCM can faithfully recover known model parameter values from synthetic rs-fMRI data. Furthermore, we assessed the utility of rDCM in application to an empirical rs-fMRI dataset acquired by the B-SNIP-1 consortium (Tamminga et al., 2013). These analyses suggest that rDCM yields biologically plausible results with regard to the functional integration during the resting state. Importantly, effective connectivity results obtained using rDCM in small networks (up to 15 nodes) were highly consistent with the estimates from spectral DCM when fitted to the identical BOLD signal time series. Furthermore, we then demonstrated that rDCM scales gracefully to large-scale network and provides sensible whole-brain effective connectivity estimates from rs-fMRI data within minutes.

First, we conducted comprehensive simulation studies to establish the face validity of rDCM when applied to synthetic resting-state fMRI data. Specifically, we found that rDCM faithfully inferred upon the data-generating (“true”) parameter values over a wide range of different settings of data quality (i.e., SNR) and sampling rate (i.e., TR). While model parameter recovery of rDCM improved for higher SNR and faster TR, consistent with previous observations in the context of task-based fMRI (Frässle et al., 2018a; Frässle et al., 2017), performance was reasonable even for the most challenging scenario (i.e., SNR = 0.5, TR = 2s). Besides this dependence on SNR and TR, model parameter recovery performance also depended on the complexity of the network with more accurate results being observed as the number of free parameters decreased.

Second, we applied rDCM to an empirical rs-fMRI dataset to study the effective connectivity amongst entire resting state networks (4 modes), as well as between their subcomponents (15 nodes). For both network architectures, results were biologically plausible and revealed inhibitory afferent and efferent connections for (regions of) the DMN, whereas (regions of) the task-positive networks elicited predominantly excitatory coupling amongst each other. This was also represented by the fact that the different intrinsic RSNs followed a hierarchical ordering (under the given metric of hierarchy strength utilized in the present study), where the DMN was situated at the bottom (primarily serving as a sink) and the task-positive networks (DAN, SAN, and CEN) were situated higher in the hierarchy (primarily serving as drivers). These findings were consistent with previous work on the hierarchical organization of RSNs (Sridharan et al., 2008; Zhou et al., 2018).

It is worth highlighting that several other studies have already applied DCM to resting-state fMRI data and have found results consistent with observations reported here. Specifically, these studies have also demonstrated a tight functional coupling amongst regions within a given RSN (Razi et al., 2015; Zhou et al., 2018) as well as comparable patterns of integration between the RSNs (Zhou et al., 2018). This consistency is noteworthy given the substantial differences in analysis strategies among these studies. First, the generative model utilized ranged from regression DCM (present work) to spectral DCM (Razi et al., 2015; Sharaev et al., 2016; Zhou et al., 2018), stochastic DCM (Li et al., 2012), and even deterministic DCM with a discrete cosine set (DCT) input (Di and Biswal, 2014). Second, studies differed in the exact choices made regarding preprocessing, the choice of regions/networks, and group level analyses. Despite these methodological and practical differences, all studies (including the present work) revealed tight coupling in the DMN between left angular gyrus and PCC, bilateral angular gyrus and the aMPFC, as well as reciprocal connections between left and right angular gyrus. In particular, the consistent involvement of the bilateral angular gyrus has been suggested to resemble its functional role in domain-general automatic processing, where the angular gyrus has been claimed to function as an automatic bottom-up buffer of incoming information (Humphreys and Lambon Ralph, 2017). Concerning functional integration amongst the subcomponents of the task-positive networks, DCM studies have so far mainly focused on task-based fMRI data. Nevertheless, these studies are still consistent with our findings in that pronounced integration was observed for regions of the SAN (Ham et al., 2013; Lamichhane and Dhamala, 2015) and DAN (Vossel et al., 2012).

Importantly, rDCM estimates were not only biologically plausible and consistent with previous findings in the literature, but also quantitatively similar to the results obtained using spectral DCM (Friston et al., 2014a). This is reassuring because spectral DCM represents a well-established variant of DCM for resting-state fMRI which has previously been tested in terms face validity (Friston et al., 2014a), construct validity (Razi et al., 2015), and test-retest reliability (Almgren et al., 2018). Hence, spectral DCM here served as a useful benchmark for assessing the construct validity of rDCM.

While group-level estimates were highly consistent across the two variants of DCM, results were more variable at the individual level. Specifically, while individual parameter estimates were highly consistent for some connections, they were more variable between the two methods for others (**Fig. 4E** and **Fig. 5F**). An interesting question for future work will be to disentangle whether the observed variability is merely due to chance or whether differences between the two methods for some of the connections actually represents meaningful differences in the variance that is picked up by rDCM and spectral DCM, respectively.

We would like to emphasize that, although rDCM and spectral DCM yielded comparable results for small networks (especially at the group level), rDCM was computationally much more efficient. This was indicated by run-times that were up to 3 orders of magnitude higher for spectral DCM for the 15-node network. While this increase in computational efficiency comes at the cost of physiological realism (e.g., fixed HRF, mean-field approximation across regions), this allows rDCM to scale gracefully to very large networks with hundreds of brain regions. In contrast, the current implementation of spectral DCM is unlikely to scale to whole-brain networks of this size, considering the run-times reported here for the small networks as well as those reported previously, suggesting that model inversion of a single DCM with 36 regions takes between 21-42h (Razi et al., 2017).

Along these lines, we here conducted an additional exploratory analysis to demonstrate the practical feasibility of whole-brain effective connectivity analyses during the resting state with rDCM. These analyses suggested that rDCM can indeed provide biologically plausible effective connectivity patterns at the whole-brain level within minutes. Specifically, rDCM revealed an expected modular structure of the effective connectivity patterns, where areas of the same lobe grouped together and showed more pronounced within-lobe interactions as compared to between-lobe interactions. This is consistent with our findings on the coupling between components of the RSNs (“nodes”) as well as previous reports on functional integration during the resting state (Fornito et al., 2016). Furthermore, (absolute) endogenous connection strengths inferred using rDCM followed a log-normal distribution, consistent with previous reports on anatomical connection strengths in mice (Oh et al., 2014; Wang et al., 2012b) and macaques (Markov et al., 2013; 2014a).

The combination of computational efficiency and applicability to simple conditions like the “resting state” renders rDCM particularly promising for application in the fields of Computational Psychiatry and Computational Neurology. Here, computational readouts of directed connectivity in whole-brain networks are of major relevance because global dysconnectivity has been postulated as a hallmark of most psychiatric and neurological disorders (Deco and Kringelbach, 2014; Fornito et al., 2015; Frässle et al., 2018b; Menon, 2011; Stephan et al., 2015). The ability to infer whole-brain effective connectivity patterns from rs-fMRI measurements is appealing from a clinical perspective as it puts minimal burden on the patient in the MR scanner. Besides these promises for clinical applications, computational efficiency and application to rs-fMRI renders rDCM also a promising tool for analyses of large-scale datasets (Sudlow et al., 2015; Van Essen et al., 2013). These datasets typically focus on rs-fMRI for its advantage in terms of easy harmonization of multi-site acquisition efforts. The vast amount of data available from these consortia calls for computational tools that are highly efficient. We anticipate that the application of rDCM to these open-source datasets may represent an exciting new opportunity to move human connectomics and network neuroscience towards directed measures of functional inegration.

It is worth mentioning that, while we replicated previous findings on putative anticorrelations between the task-negative network (DMN) and the task-positive networks (DAN, SAN) during the resting state (Fox et al., 2005), one should be cautious with regard to the interpretation of this finding. Specifically, it has been argued that the observed anticorrelations may – at least in part – be a consequence of specific preprocessing choices. In particular, global signal regression has been argued to artificially induce anticorrelations by removing any positive correlation between DMN and DAN/SAN that may be hidden in the signal (Buckner et al., 2008; Fox et al., 2009; Murphy et al., 2009). In the present work, we did perform global signal regression to correct for potential confounds like psychological, hardware, and motion artifacts. Hence, results obtained with rDCM in the present study are consistent with previous reports that followed comparable preprocessing strategies (e.g., global signal regression). However, our methodological study cannot – and does not intend – to resolve the ongoing controversy on the validity of anticorrelations between RSNs.

Finally, we would like to highlight that the development of rDCM is still in an early stage and the present implementation is subject to several limitations. First, in its current form, rDCM does not account for variability in the hemodynamic response across regions and subjects (Aguirre et al., 1998; Handwerker et al., 2004). Hence, a major future development is to replace the fixed HRF with a more flexible hemodynamic model that nevertheless preserves the computational efficiency of rDCM. Second, it is possible that explicitly accounting for endogenous neuronal fluctuations in the generative forward model could further increase the explanatory power of rDCM for rs-fMRI data. Third, the linear formulation of rDCM and the stationarity assumption of the Fourier transform implies that only stationary measures of functional integration can be obtained. However, functional integration changes dynamically on short timescales, under the influence of synaptic plasticity and neuromodulation (Stephan et al., 2015). Hence, another important future development is to introduce dynamics back into the effective connectivity estimates obtained with rDCM.

Despite these limitations, our results demonstrate the face and construct validity of rDCM for modeling rs-fMRI data: they demonstrate good recovery of known connection strengths from simulated data, illustrate that rDCM can infer biologically plausible effective connectivity patterns from empirical rs-fMRI data and show that rDCM estimates are consistent with results from spectral DCM, a well-established DCM variant for rs-fMRI. Importantly, due to the computational efficiency of our method, rDCM scales gracefully with network size and provides sensible whole-brain resting-state effective connectivity patterns. This opens up promising new avenues for whole-brain connectomics based on directed measures of functional integration during the resting state. Such a readout of whole-brain effective connectivity will hopefully not only provide novel insights into the organizing principles of the human brain in health, but might also prove valuable for unraveling pathophysiological mechanisms and for enabling single-subject predictions about clinical outcomes (Frässle et al., 2020b).

## 5 Code and Data Availability

A Matlab implementation of the regression dynamic causal modeling (rDCM) approach is available as open source code in the **T**ranslational **A**lgorithms for **P**sychiatry-**A**dvancing **S**cience (TAPAS) toolbox (www.translationalneuromodeling.org/software). Extensions to the rDCM toolbox that facilitate its application to rs-fMRI data will be made available as part of the next release. Furthermore, following acceptance of this paper, we will publish the code for the analysis online as part of an online repository that conforms to the FAIR (Findable, Accessible, Interoperable, and Re-usable) data principles. The fMRI data were acquired by the B-SNIP-1 consortium and access to the data is possible upon reasonable request.

## 6 Acknowledgements

This work was supported by the UZH Forschungskredit Postdoc (SF), the ETH Zurich Postdoctoral Fellowship Program and the Marie Curie Actions for People COFUND Program (SF), the Strategic Focal Area “Personalized Health and Related Technologies (PHRT)” of the ETH Domain grant #2017-403 (SJH), the René and Susanne Braginsky Foundation (KES), the Swiss National Science Foundation 320030_179377 (KES), and the University of Zurich (KES).

## Supplementary Material

## Supplementary Material S1: Details on Data Acquisition

Structural and functional MRI data were acquired at the five sites of the Bipolar-Schizophrenia Network on Intermediate Phenotypes (B-SNIP-1) consortium (Tamminga et al., 2013; 2014). These included the University of Maryland School of Medicine (Baltimore), Commonwealth Research Center at Harvard Medical School (Boston), University of Chicago Medical Center (Chicago), University of Texas Southwestern Medical Center (Dallas), and Olin Neuropsychiatry Research Center at the Institute of Living (Hartford). Participants were scanned on 3-Tesla MR scanners of different manufacturers, including GE Signa, Siemens Trio, Philips Achieva, and Siemens Allegra. For the present methodological study, neuroimaging data from the sites in Baltimore, Chicago, Dallas and Hartford was utilized since not all information relevant for our preprocessing pipeline was available for the Boston data.

For *Baltimore*, resting-state fMRI data was acquired on a 3T Siemens Tim Trio scanner. Volumes were acquired using a T_2_^*^-weighted echo-planar imaging (EPI) sequence (36 slices, TR=2210ms, TE=30ms, field of view (FoV) 224×224×144mm^3^, voxel size 3.5×3.5×4mm^3^, flip angle 70°). In addition to the functional images, a high-resolution anatomical image was acquired for each participant using a T1-weighted Magnetization Prepared Rapid Gradient Echo (MPRAGE) sequence, following the Alzheimer’s Disease Neuroimaging Initiative protocol (http://adni.loni.usc.edu). The MPRAGE sequence parameters were as follows: 3D acquisition, sagittal slab, 160 slices, shot interval 2300ms, inversion time 900ms, TR=7.2ms, TE=2.91ms, field of view (FoV) 256×240×192mm^3^, voxel size 1×1×1.2mm^3^, flip angle 9°.

For *Chicago*, resting-state fMRI data was acquired on a 3T GE Signa HDx scanner. Volumes were acquired using a T_2_^*^-weighted echo-planar imaging (EPI) sequence (29 slices, TR=1775ms, TE=27ms, field of view (FoV) 224×224×145mm^3^, voxel size 3.5×3.5×4mm^3^, slice gap 1mm, flip angle 60°). A high-resolution anatomical image was acquired using a T1-weighted MPRAGE sequence (3D acquisition, sagittal slab, 166 slices, shot interval 2300ms, inversion time 700ms, TR=6.988ms, TE=2.848ms, field of view (FoV) 260×260×200mm^3^, voxel size 1.016×1.016×1.2mm^3^, flip angle 8°).

For *Dallas*, resting-state fMRI data was acquired on a 3T Philips Achieva scanner. Volumes were acquired using a T_2_^*^-weighted echo-planar imaging (EPI) sequence (29 slices, TR=1500ms, TE=27ms, field of view (FoV) 224×224×145mm^3^, voxel size 3.5×3.5×4mm^3^, slice gap 1mm, flip angle 60°). A high-resolution anatomical image was acquired using a T1-weighted MPRAGE sequence (3D acquisition, sagittal slab, 170 slices, shot interval 3000ms, inversion time 846ms, TR=6.8ms, TE=3.1ms, field of view (FoV) 256×256×204mm^3^, voxel size 1×1×1.2mm^3^, flip angle 8°).

And finally, for *Hartford*, resting-state fMRI data was acquired on a 3T Siemens Allegra scanner. Volumes were acquired using a T_2_^*^-weighted echo-planar imaging (EPI) sequence (29 slices, TR=1500ms, TE=27ms, field of view (FoV) 224×224×145mm^3^, voxel size 3.5×3.5×4mm^3^, slice gap 1mm, flip angle 70°). A high-resolution anatomical image was acquired using a T1-weighted MPRAGE sequence (3D acquisition, sagittal slab, 160 slices, shot interval 2300ms, inversion time 900ms, TR=6.8ms, TE=2.91ms, field of view (FoV) 256×240×192mm^3^, voxel size 1×1×1.2mm^3^, flip angle 9°).

## Supplementary Figures

**Supplementary Figure S1.**
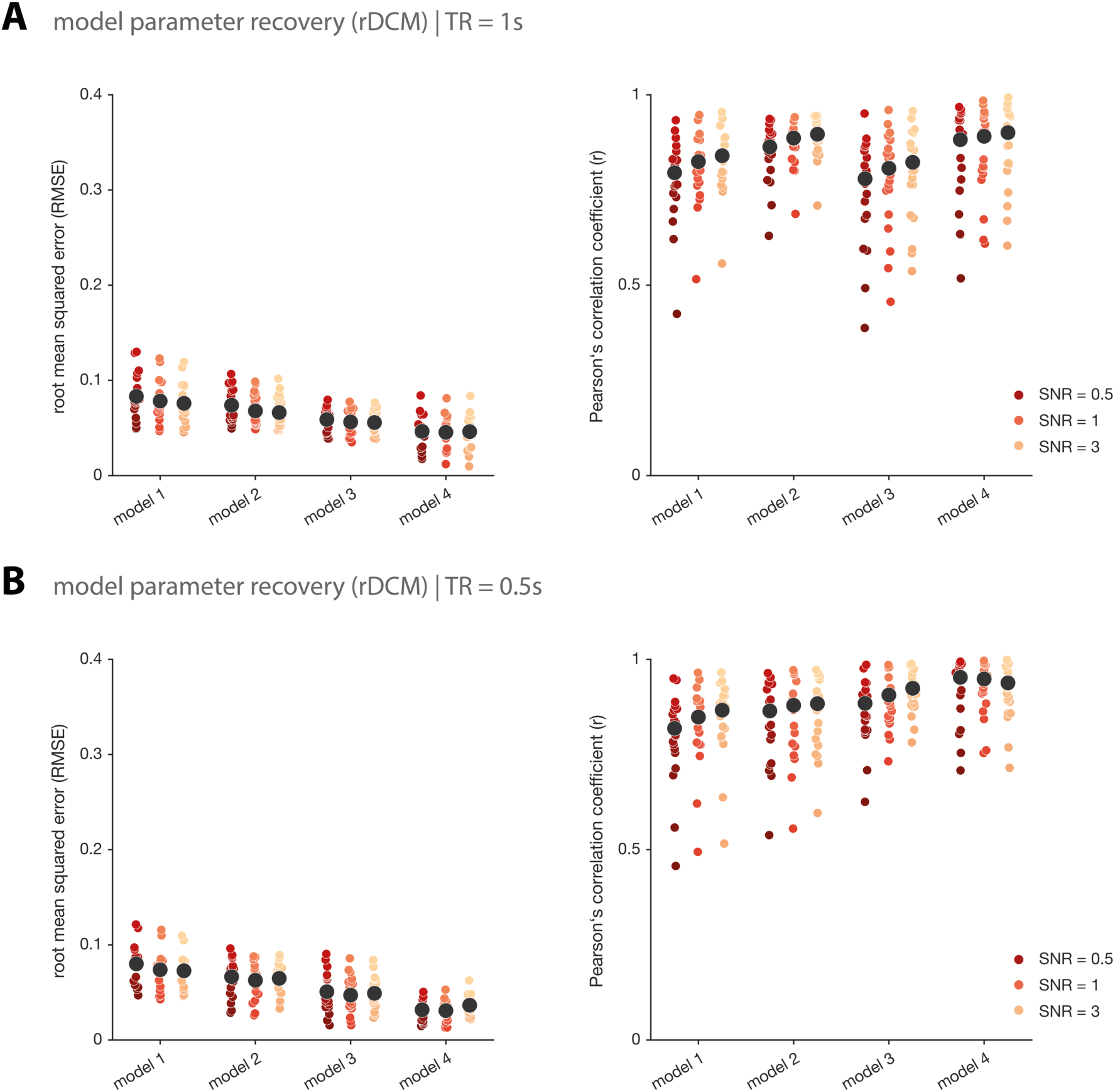
Model parameter recovery for regression DCM. Model parameter recovery for synthetic resting-state fMRI data in terms of the root mean squared error (*left*) and the Pearson’s correlation coefficient (*right*) between the inferred and the “true” data-generating parameter values for regression DCM (rDCM) for **(A)** a repetition time (TR) of 1s, and **(B)** a repetition time (TR) of 0.5s. Each colored dot represents the respective RMSE or Pearson’s correlation coefficient for an individual synthetic subject. Black dots represent the mean RMSE or mean Pearson’s correlation coefficient, averaged across all 20 synthetic subjects. Note that averaging of the individual Pearson’s correlation coefficients was performed in z-space (see main text in main manuscript). Results are shown for four different network architectures, where model 1 is the most complex (full) model and model 4 is the sparsest network. Furthermore, results are shown for an exemplary repetition time (TR) of 2s and three different signal-to-noise ratios (SNRs): 0.5 (red), 1 (orange), and 3 (yellow).

**Supplementary Figure S2.**
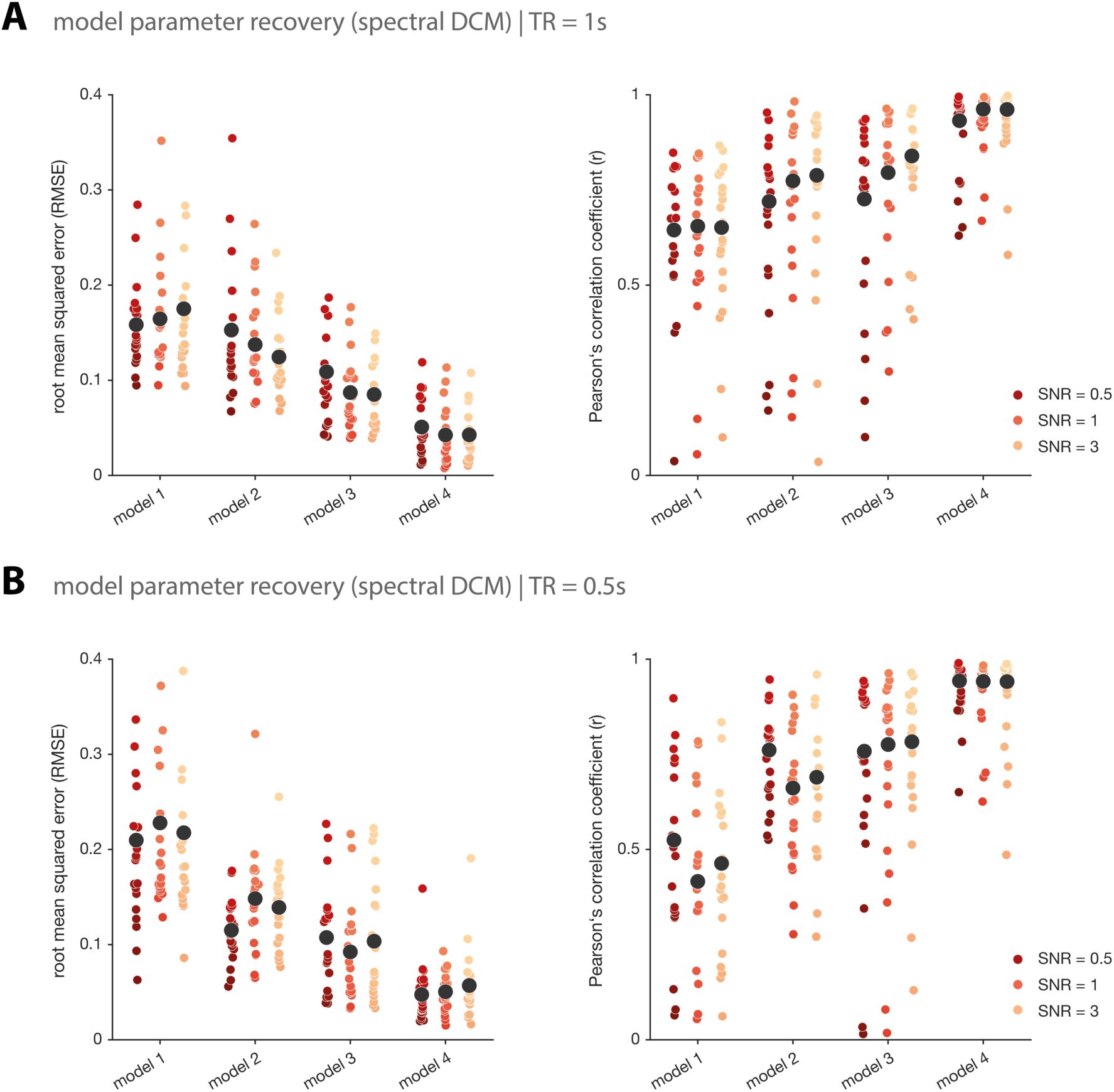
Model parameter recovery for spectral DCM. Model parameter recovery for synthetic resting-state fMRI data in terms of the root mean squared error (*left*) and the Pearson’s correlation coefficient (*right*) between the inferred and the “true” data-generating parameter values for spectral DCM for **(A)** a repetition time (TR) of 1s, and **(B)** a repetition time (TR) of 0.5s. Each colored dot represents the respective RMSE or Pearson’s correlation coefficient for an individual synthetic subject. Black dots represent the mean RMSE or mean Pearson’s correlation coefficient, averaged across all 20 synthetic subjects. Note that averaging of the individual Pearson’s correlation coefficients was performed in z-space (see main text in main manuscript). Results are shown for four different network architectures, where model 1 is the most complex (full) model and model 4 is the sparsest network. Furthermore, results are shown for an exemplary repetition time (TR) of 2s and three different signal-to-noise ratios (SNRs): 0.5 (red), 1 (orange), and 3 (yellow).

## Footnotes

1 In the Brainnetome nomenclature, this set of regions is called “LIM” (limbic). However, as the term “limbic” is not well-defined (Kötter & Stephan, 1997) and since “LIM” exclusively consists of cingulate areas, we prefer to call this set of regions “CNG” (cingulate).

